# Colistin susceptibility pattern in gram negative bacilli isolated from patients of dhaka medical college hospital with distribution of antibiotic resistance genes among them

**DOI:** 10.1101/2020.04.16.045906

**Authors:** Hasbi Ara Mostofa, S.M. Shamsuzzaman, MD. Maniul Hasan

## Abstract

Colistin is one of the last resort of antibiotics used specifically in the treatment of multidrug resistant gram negative bacilli. Though an older class of antibiotic with no use in last four decades, resurgence of colistin use is also causing emergence of colistin resistant gram negative bacilli.The aim of this cross sectional study was to investigate the colistin susceptibility and resistant pattern along with colistin resistant genes in gram negative bacilli among different samples in Dhaka medical college hospital. Total 300 samples (wound swab, urine, endotracheal aspirate, blood and sputum) were collected from 2015 to 2016. 204 gram negative bacilli were isolated and tested for resistance to colistin by disc diffusion method. Then, the prevalence of colistin resistant genes (pmrA, pmrB, pmrC, mgrB, phoP, phoQ, lpxA, lpxC, lpxD, mcr-1) was detected by polymerase chain reaction method (PCR). Among 204 gram negative bacilli, 22 (10.78%) colistin resistance was identified. PCR analysis revealed that 50% of these isolates contained pmrA gene, followed by phoP gene (27.27%). The prevalence of pmrB, phoQ, mgrB, lpxC and lpxD were 22.72%, 13.63%, 13.63%, 13.63%, 13.63%, and 13.63% respectively among the isolates. Colistin resistance is widespread among gram negative bacilli isolated from human infections. Colistin resistance (10.78%) found in the study is quite alarming. A restricted and rational use of the colistin is the need of hour.

## 1. Introduction

Colistin was first isolated in Japan by Koyama and coworker from the spore forming soil bacterium Bacillus polymyxa subsp. colistinus in 1947 and first used as an intravenous formulation in the 1950s (1). Colistin has been approved by the US FDA and has been available since 1959 for the treatment of infections caused by gram negative bacteria because these agents are rapidly bactericidal against them (2).

Resistance to colistin is mediated mainly through lipid A structural adjustments, resulting from the addition of phosphoethanolamine (Pet) and 4-amino-4-deoxy-L-arabinose (L-Ara-4N) to the lipid A moiety on the surface membrane; these additions make lipid A less cationic such that the anionic colistin is unable to bind and initiate membrane lysis (3,4). Genes that encode enzymes involved in the synthesis of lipid A are the pmrHFIJFKLM (also known as arnBCADTEFpmrE) in Gram-negative bacteria in addition to lpxA, lpxC and lpxD in *Acinetobacter baumannii*. These genes are in turn regulated by pmrAB, phoPQ and mgrB genes. Hence, mutations in any of these genes result in a defect in lipid A synthesis and/or addition of L-Ara-4N or PEt to lipid A, leading to colistin resistance through reduced anionic charges.

Recently, a plasmid-borne PET transferase, mcr-1, has been identified in China, Malaysia and Laos, in swine, raw meat and hospitalized patients (5). This study was conducted to observe the distribution of colistin resistance genes among the gram negative bacilli isolated from patients of Dhaka Medical College Hospital (DMCH).

## 3. Methods

### 3.1. Isolation and Identification of gram negative bacilli

#### Ethical issue

After getting ethical assurance from ethical review committee of DMCH and informed written consent was observed from the patients admitted in DMCH.

A total of 204 gram negative bacilli isolates from wound swab, urine, endotracheal aspirate, blood and sputum samples were obtained from patient of Dhaka medical college hospital, Dhaka, Bangladesh from July 2015 to June 2016. All the samples were streaked on to Blood agar media and MacConkey agar and incubated at 37°C for 24 hours. Then, the plates were examined for gram negative bacilli. All the organisms were identified by colony morphology, hemolytic criteria, staining character, pigment production and biochemical tests as per standard technique (6). The isolated organisms were inoculated in TSI, MIU, citrate agar media and organisms were identified based on biochemical reactions (6).

### 3.2. Antibiotic Susceptibility Testing

colistin susceptibility to antimicrobial agents of all isolates was done by Kirby Bauer modified disk diffusion technique using Mueller Hinton agar plates and zones of inhibition were interpreted according to guidelines (CLSI, 2015; EUCAST, 2015) Antibiotic disc colistin (10 µg) was used. Inoculums were prepared and the turbidity of the suspension was compared with McFarland turbidity standard 0.5 (6). The swab was streaked evenly over the surface of Mueller Hinton agar plate in three directions rotating the plate approximately 60°C. The inoculating plate was then allowed to dry for 3-5 minutes. Antibiotic discs were placed on the inoculating plate 15 mm away from the edge of the plate and 25 mm apart from one disc to another from centre to centre (6). The inoculated plates were incubated aerobically at 37°C overnight. After overnight incubation, The zone of inhibition were evaluated with standard chart of CLSI guideline (2015) for *Pseudomonas aeruginosa* and *Acinetobacter baumannii*. EUCAST guideline was followed for colistin disc in case of *Enterobacteriaceae*.

### 3.3. Identification of colistin resistant Genes using PCR Assay

PCR amplification assays were performed to determine the prevalence of pmrA, pmrB, pmrC, mgrB, lpxA, lpxC, lpxD, phoP, phoQ, and mcr-1 among all colistin resistant gram negative bacilli. Table 1 demonstrates the sets of primers used to identify these determinants. DNA was extracted by boiling method. Three hundred μl of sterile distilled water was added into microcentrifuge tubes having bacterial pellet and vortexed until mixed. Mixture was heated at 100°C for 10 minutes in a heat block. After heating, immediately the microcentrifuge tubes were placed on ice for 5 minutes and then centrifuged at 14,000 rpm at 4°C for 6 minutes. Supernatant was taken into another microcentrifuge tube by micropipette and was used for PCR. Extracted DNA was preserved at 4°C for 7-10 days and - 20°C for long time.Then primers were mixed with Tris-EDTA (TE) buffer according to manufacturer’s instruction. For each sample, a total 25μl of mixture was prepared by mixing of 12.5μl of mastermix (mixture of dNTP, taq polymerase Mgcl2 and PCR buffer), 2 μl of forward primer, 2μl of reverse primer (promega corporation, USA), 2μl of DNA template and 6.5μl of sterile distilled water in a PCR tube. After a brief vortex, the tubes were centrifuged in a microcentrifuge for few seconds.

**Table 1.**
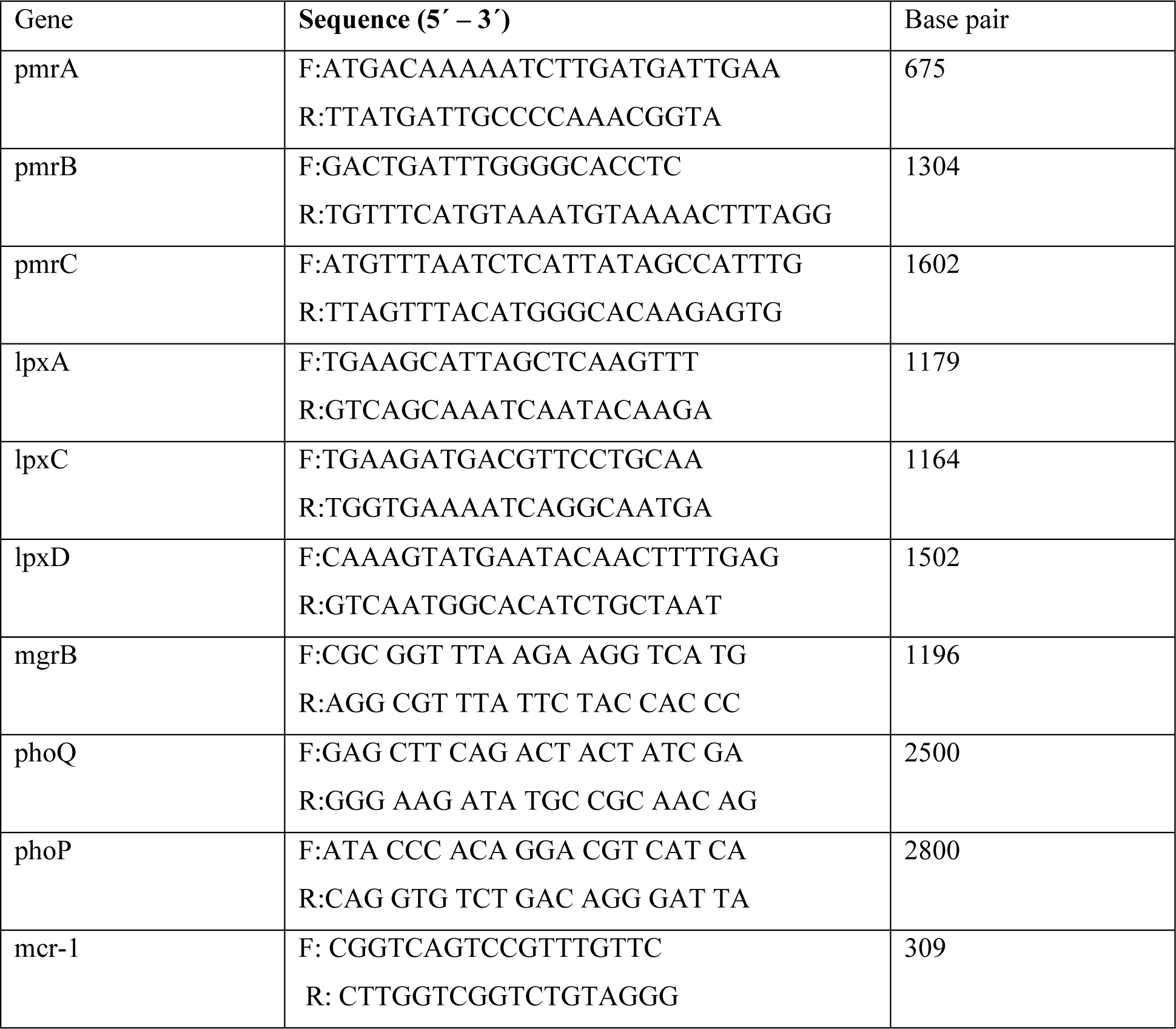
Primer Sequences Used for PCR Identification of pmrA, pmrB, pmrC, mgrB, lpxA, lpxC, lpxD, phoP, phoQ and mcr-1..

PCR reaction consisted of initial denaturation at 95°C for 10 minutes, then 30 cycles of denaturation at 95°C for one minute, annealing for 45 seconds, extention at 72°C for one minute and 30 seconds and final extention at 72°C for 10 minutes. Then the amplicons were kept at 4°C until gel electrophoresis. Annealing temperature varies with GC contents of primers. Finally, the reaction products were run on 1% agarose gel, and the prevalence of each colistin resistant gene was determined using agarose gel electrophoresis. The size of the amplified DNA was assessed comparing with the bands of DNA ladder (Figure1 and Figure 2).

**Figure 1:**
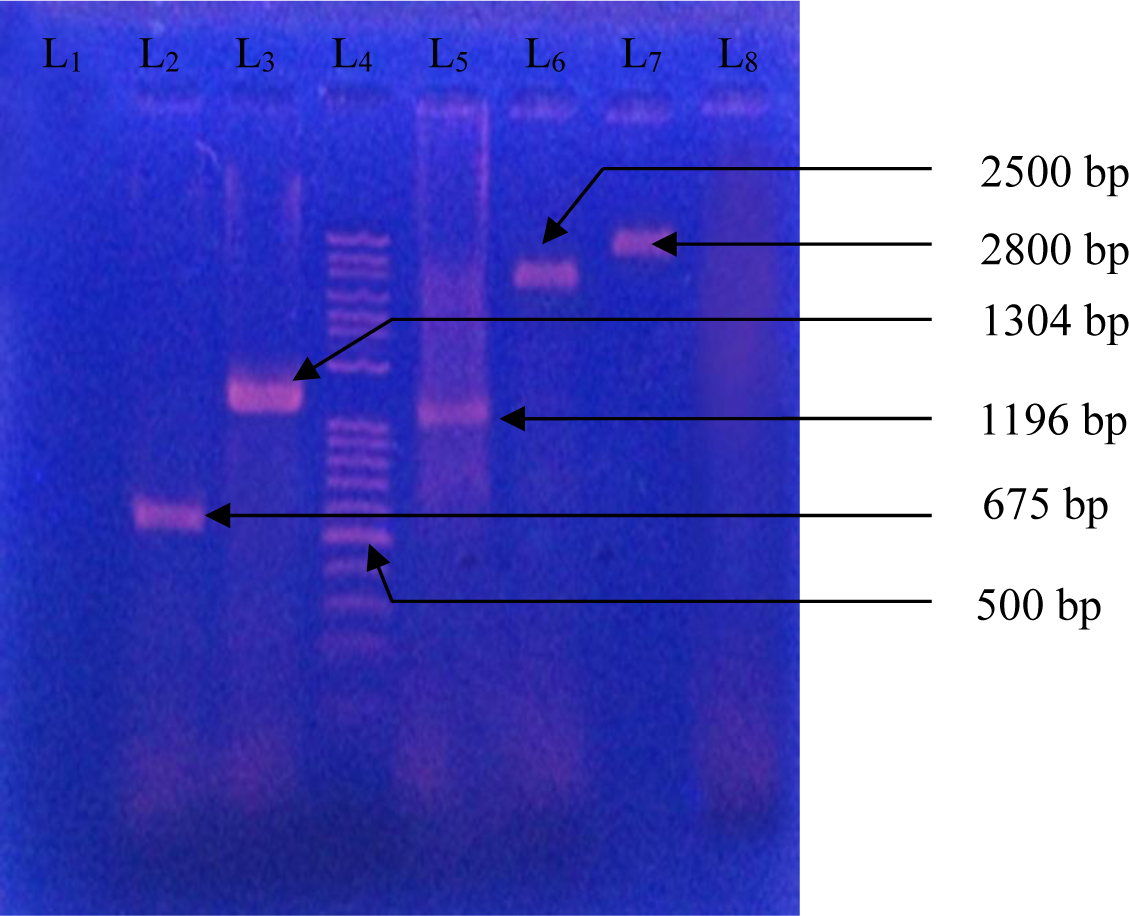
Photograph of gel electrophoresis of amplified DNA of 675 bp for pmrA gene (lane 2), DNA of 1304 bp for pmrB gene (lane 3), one kb base pair DNA ladder (lane 4), DNA of 1196 bp for mgrB gene (lane 5), DNA of 2500 bp for phoP gene (lane 6), DNA of 2800 bp for phoQ gene (lane 7), Negative control *Escherichia coli* ATCC 25922 (lane 8).

**Figure 2:**
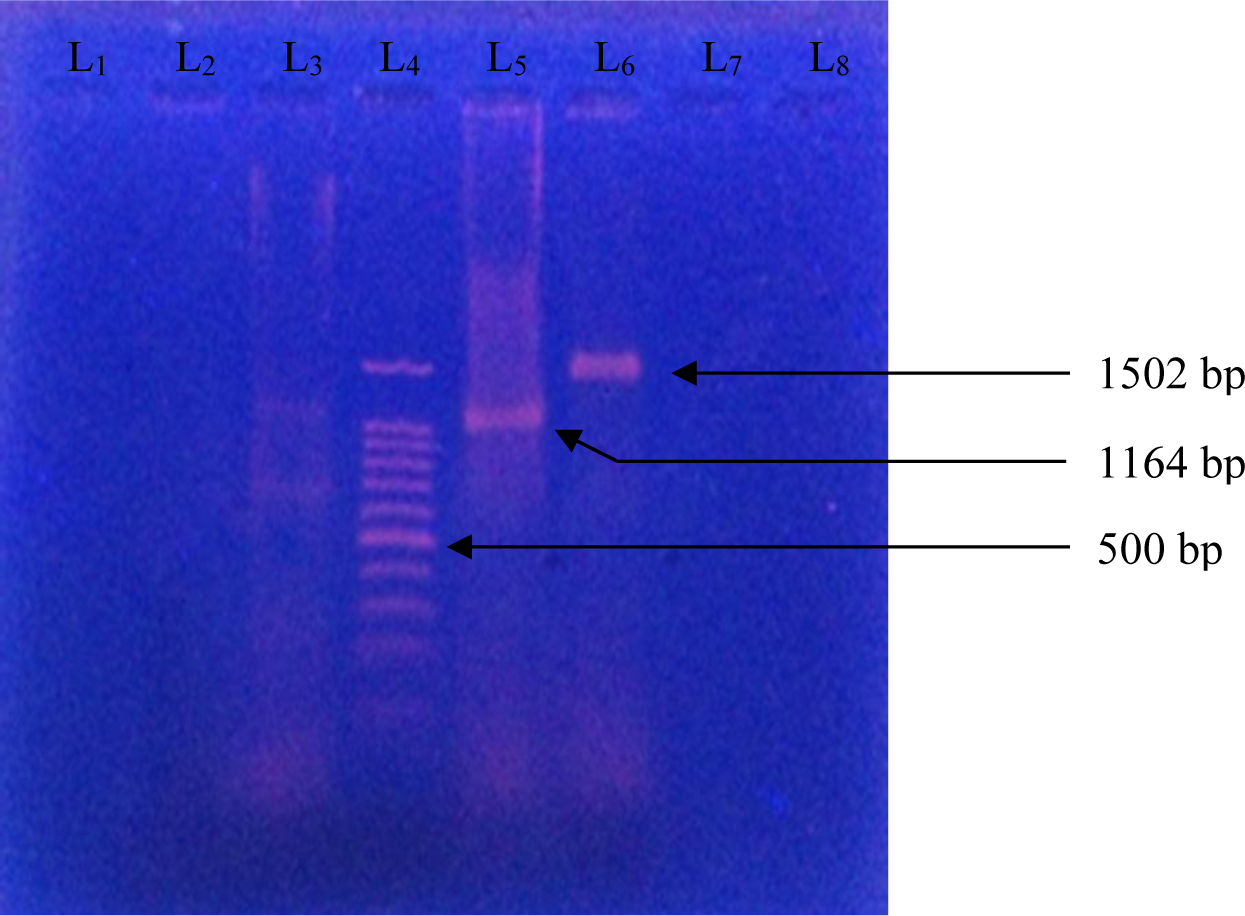
Photograph of gel electrophoresis of amplified DNA of 1164 bp for *lpxC* gene (Lane 5), DNA of 1502 bp for lpxD gene (Lane 6), one kb DNA ladder (Lane 4), Negative control without DNA (Lane 3).

### 3.4 DNA sequencing

DNA sequencing of pmrA, pmrB, mgrB, lpxC, lpxD, phoP and phoQ genes of colistin were done. For sequencing of bacterial DNA, purification of amplified PCR product were done by following (FAVOGEN, Biotech Corp.).

## 4. Results

The results of antibiotic susceptibility testing in gram negative bacillii showed that 22 (10.78%) were phenotypically and genotypically resistant to colistin, (89.22%) were susceptible to colistin (Table 2). The frequency and distribution of colistin resistance genes among gram negative bacilli in the present study are demonstrated in (Table 3). The results of the present study revealed that pmrA was the most prevalent gene detected in 50% of the colistin resistant isolates, followed by phoP (27.27%), and pmrB (22.72%). Moreover, phoQ, mgrB, lpxC and lpxD each were found in 13.63% and these were the lowest prevalent gene among the studied determinants. pmrA (*Escherichia coli*) gene in this present study had mutation at 369 position the amino acid leucine is replaced by valine. pmrA (*Pseudomonas aeruginosa*) gene in the present study had mutation at 7 position where the amino acid arginine is replaced by leucine. pmrB (*Pseudomonas aeruginosa*) gene in the present study had mutation at 14 position where the amino acid proline is replaced by leucine. PhoP (*Pseudomonas aeruginosa*) gene in the present study had mutation at 237 position where the amino acid leucine is replaced by arginine. phoP (*Proteus mirabilis)* gene in the present study had mutation at 31 position where the amino acid glutamic acid is replaced by glutamin, at 38 position where serine is replaced by histidine, at 43 position where glycine is replaced by serine, at 193 position leucine is replaced by isoleucine and at 209 position where serine is replaced by threonine. phoQ (*Pseudomonas aeruginosa*) gene in the present study had mutation at 205 position in which phenylalanine is replaced by tyrosine. lpxC (*Acinetobacter baumannii*) gene in the present study had mutation at 223 position where the amino acid lysine is replaced by serine, at 224 position where aspartic acid is replaced by glutamine. lpxD (*Acinetobacter baumannii*) gene in the present study had mutation at 50 position where the amino acid valine is replaced by isoleucine at 371 position where isoleucine is replaced by valine. MgrB (*Klebsiella pneumoniae*) gene in the present study had mutation at 35 position where the amino acid methionine is replaced by valine and deletion in one amino acid at 83 position. But it is not clear whether changed protein is associated with resistance to colistin. In the present study, mcr-1, lpxA and pmrC were extensively searched but none other isolates were found positive for any of the genes.

**Table 2:** Colistin resistant organisms among different species of isolated gram negative bacteria from different samples. (N=204)

| Organism | Wound Swab<br>n(%) | Urine<br>n(%) | ETA<br>n(%) | Blood<br>n(%) | Sputum<br>n(%) | Total<br>n(%) |
| --- | --- | --- | --- | --- | --- | --- |
| *Esch.coli*<br>(N=43) | 1(2.32) | 0(0.00) | 0(0.00) | 0(0.00) | 0(0.00) | 1(2.32) |
| *K. pneumoniae*<br>(N=26) | 1(3.84) | 0(0.00) | 0(0.00) | 0(0.00) | 0(0.00) | 1(3.84) |
| *K. oxytoca*<br>(N=13) | 1(7.69) | 0(0.00) | 0(0.00) | 0(0.00) | 0(0.00) | 1(7.69) |
| *C. freundii*<br>(N=16) | 0(0.00) | 0(0.00) | 1(6.25) | 0(0.00) | 0(0.00) | 1(6.25) |
| *C. koseri*<br>(N=5) | 0(0.00) | 0(0.00) | 0(0.00) | 0(0.00) | 0(0.00) | 0(0.00) |
| *P. vulgaris*<br>(N=7) | 1(14.28) | 0(0.00) | 0(0.00) | 0(0.00) | 0(0.00) | 1(14.28) |
| *P. mirabilis*<br>(N=6) | 1(16.7) | 0(0.00) | 0(0.00) | 0(0.00) | 0(0.00) | 1(16.70) |
| *E. aerogenes*<br>(N=10) | 0(0.00) | 0(0.00) | 0(0.00) | 0(0.00) | 0(0.00) | 0(0.00) |
| *Salmonella*<br>*spp*(N=2) | 0(0.00) | 0(0.00) | 0(0.00) | 0(0.00) | 0(0.00) | 0(0.00) |
| *A. baumannii*<br>(N=24) | 0(0.00) | 0(0.00) | 3(12.50) | 0(0.00) | 0(0.00) | 3(12.50) |
| *P.aeruginosa*<br>(N=52) | 7(13.46) | 4(7.69) | 2(3.84) | 0(0.00) | 0(0.00) | 13(25.00) |
| Total | 12(5.90) | 4(1.96) | 6(2.94) | 0(0.00) | 0(0.00) | 22(10.78) |

**Table 3.** The frequency of colistin resistant Genes in gram negative bacilli.

| organism | pmrA(%) | pmrB(%) | phoP (%) | phoQ(%) | mgrB(%) | lpxC(%) | lpxD(%) |
| --- | --- | --- | --- | --- | --- | --- | --- |
| *Esch. Coli*<br>(N=1) | 1(100) | 0(0.00) | 0(0.00) | 0(0.00) | 0(0.00) | 0(0.00) | 0(0.00) |
| *Klebsiella*<br>*pneumoniae*<br>(N=1) | 0(0.00) | 0(0.00) | 0(0.00) | 0(0.00) | 1(100) | 0(0.00) | 0(0.00) |
| *Klebsiella*<br>*oxytoca*<br>(N=1) | 0(0.00) | 0(0.00) | 0(0.00) | 0(0.00) | 1(100) | 0(0.00) | 0(0.00) |
| *Citrobacter*<br>*freundii*<br>(N=1) | 1(100) | 0(0.00) | 0(0.00) | 0(0.00) | 1(100) | 0(0.00) | 0(0.00) |
| *Proteus*<br>*vulgaris*<br>(N =1) | 1(100) | 0(0.00) | 0(0.00) | 0(0.00) | 0(0.00) | 0(0.00) | 0(0.00) |
| *Proteus*<br>*mirabilis*<br>(N=1) | 0(0.00) | 0(0.00) | 1(100) | 0(0.00) | 0(0.00) | 0(0.00) | 0(0.00) |
| *A.baumannii*<br>(N=3) | 0(0.00) | 0(0.00) | 0(0.00) | 0(0.00) | 0(0.00) | 3(100) | 3(100) |
| *P.aeruginosa*<br>(N=13) | 8(61.53) | 5(38.5) | 5(38.46) | 3(23.07) | 0(0.00) | 0(0.00) | 0(0.00) |
| Total | 11(50.00) | 5(22.72) | 6(27.27) | 3(13.63) | 3(13.63) | 3(13.63) | 3(13.63) |

## 5. Discussion

Among 204 gram negative bacilli isolated from wound swab, urine, endotracheal aspirate, blood, and sputum samples, 22 strains showed resistance to colistin. Monitoring the distribution prevalence of colistin resistance genes, we found that the pmrA gene had the highest frequency among the studied determinants. In this present study, prevalence of colistin resistant gram negative bacilli was 10.78%. Percentage of colistin resistance ranged from (1.9% - 3.3%) in 2007, 8% of the *Pseudomonas aeruginosa*, 12% of resistance among *Acinetobacter baumannii*, 4.65% in 2014, 9.98% in 2016 (13, 14, 15, 19, 20). In Bangladesh antibiotic resistance increased due to indiscriminate antibiotic use, sale of antibiotic over the counter, offer alternate antibiotic by drug seller and no drug selling policy (18).

Adaptive or mutational mechanism is the basis for development of colistin resistance in gram negative bacilli.(10,11) Mutations result in alteration in outer membrane of gram negative bacilli which is the site of colistin action (11). Also plasmid mediated colistin resistance has been established in animal and humans (16, 17).

In the present study, out of 22 colistin resistant gram negative bacilli, pmrA gene was present in 50% of the isolates (Table 3). This study observed that 61.53% colistin resistant *Pseudomonas aeruginosa* were positive for pmrA gene. A study by Lee and Ko (2014) reported that 43.75% colistin resistant *Pseudomonas aeruginosa* were positive for pmrA gene (21). Increased use of the cationic antibiotic colistin to treat multidrug-resistant *Acinetobacter baumannii, Pseudomonas aeruginosa* and *Klebsiella pneumonia* has led to the development of colistin-resistant strains and treatment of patients with colistin can induce not only increased resistance to colistin but also resistance to host cationic antimicrobials *(29).*

pmrA (*Escherichia coli*) gene in this present study had mutation at 369 position the amino acid leucine is replaced by valine. pmrA (*Pseudomonas aeruginosa*) gene in the present study had mutation at 7 position where the amino acid arginine is replaced by leucine. PmrA gene had mutation at 157 position where leucine is replaced by glutamine (21). pmrB (*Pseudomonas aeruginosa*) gene in the present study had mutation at 14 position where the amino acid proline is replaced by leucine. PmrB gene had mutation at 15, 48, 67, 70 and 167 position where amino acid valine replaced by isoleucine, methionine replaced by leucine, alanine replaced by threonine, aspartic acid replaced by asparagines, leucine is replaced by proline respectively (21). PhoP (*Pseudomonas aeruginosa*) gene in the present study had mutation at 237 position where the amino acid leucine is replaced by arginine. phoP (*Proteus mirabilis)* gene in the present study had mutation at 31 position where the amino acid glutamic acid is replaced by glutamin, at 38 position where serine is replaced by histidine, at 43 position where glycine is replaced by serine, at 193 position leucine is replaced by isoleucine and at 209 position where serine is replaced by threonine.phoP gene had mutation at 26 and 385 position where leucine is replaced by glutamine and glycin is replaced by serine respectively (4). phoQ (*Pseudomonas aeruginosa*) gene in the present study had mutation at 205 position in which phenylalanine is replaced by tyrosine. PhoQ gene had mutation at 123 and 143 position where alanine replaced by valine and lysine replaced by glutamine (21). lpxC (*Acinetobacter baumannii*) gene in the present study had mutation at 223 position where the amino acid lysine is replaced by serine, at 224 position where aspartic acid is replaced by glutamine.lpxC had mutation at 141 and 158 position where lysine replaced by arginine and serine replaced by arginine repectively (28). lpxD (*Acinetobacter baumannii*) gene in the present study had mutation at 50 position where the amino acid valine is replaced by isoleucine, at 371 position where isoleucine is replaced by valine.lpxD had mutation at 102, 104, 141, 173 and 178 position where serine replaced by threonine, valine replaced by isoleucine, arginine replaced by glycine, threonine replaced by lysine, isoleucine replaced by valine respectively (28). MgrB (*Klebsiella pneumoniae*) gene in the present study had mutation at 35 position where the amino acid methionine is replaced by valine and deletion in one amino acid at 83 position. MgrB had mutation at 48 position where stop codon replaced by threonine. But it is not clear whether changed protein is associated with resistance to colistin. In the present study, mcr-1, lpxA and pmrC were extensively searched but none other isolates were found positive for any of the genes..

The results of monitoring the prevalence of colistin resistance genes in gram negative bacilli isolated from wound swab, urine, endotracheal aspirate, blood and urine confirmed the emergence and spreading of the colistin resistant gram negative bacilli in human populations.

## Acknowledgments

The authors greatfully acknowledged by department of microbiology of Dhaka Medical College, Dhaka, Bangladesh for providing laboratory support.

